# SCC*mec* transformation requires living donor cells in mixed biofilms

**DOI:** 10.1101/2023.09.11.557198

**Authors:** Mais Maree, Yuri Ushijima, Masato Higashide, Kazuya Morikawa

## Abstract

Methicillin-resistant *Staphylococcus aureus* (MRSA) is an important human pathogen that has emerged through the horizontal acquisition of the staphylococcus cassette chromosome *mec* (SCC*mec*). Previously, we showed that SCC*mec* from heat-killed donors can be transferred via natural transformation in biofilms at frequencies of 10^-8^-10^-7^. Here, we show an improved transformation assay of SCC*mec* with frequencies up to 10^-2^ using co-cultured biofilms with living donor cells. The Ccr-attB system played an important role in SCC*mec* transfer, and the deletion of *ccrAB* recombinase genes reduced the frequency ∼30-fold. SCC*mec* could be transferred from either MRSA or methicillin-resistant coagulase-negative staphylococci to some of methicillin-sensitive *S. aureus* recipients. In addition, the transformation of other plasmid or chromosomal genes is enhanced by using living donor cells. This study emphasizes the role of natural transformation as an evolutionary ability of *S. aureus* and in MRSA emergence.

**Importance:** Methicillin-resistant *Staphylococcus aureus* (MRSA) stands out as the leading contributor to fatalities attributed to antibiotic-resistant infections. To comprehend its emergence and dissemination, it is crucial to understand the mechanisms behind it. MRSA has arisen through the horizontal acquisition of the methicillin resistance gene *mecA*, which is harboured within the staphylococcal cassette chromosome (SCC*mec*). Our study sheds light on a noteworthy discovery: when methicillin-sensitive *S. aureus* recipient cells are grown alongside viable methicillin-resistant donor cells in co-cultured biofilms, a highly efficient transfer of SCC*mec* occurs, both within the same species and between different species. This remarkable transfer process is facilitated by natural transformation, underscoring its pivotal role in the evolution of staphylococci and the emergence of MRSA.

## Introduction

*Staphylococcus aureus* is a Gram-positive bacterium that commonly inhabits the skin and mucosal surfaces of animals and humans. As an opportunistic pathogen, it causes a wide variety of diseases, ranging from minor skin abscesses and food poisoning to endocarditis, toxic shock syndrome, and sepsis. *S. aureus* is notorious for its ability to acquire antibiotic resistance (1, 2), and methicillin-resistant *S. aureus* (MRSA) has significant mortality and morbidity as the leading cause of nosocomial infections (3). MRSA has also spread to the community (community-associated MRSA, CA-MRSA) and has been associated with the livestock (livestock-associated MRSA, LA-MRSA), posing a serious health and economic burden on a global scale (4, 5).

MRSA emerges through the acquisition of a mobile genetic element known as staphylococcal cassette chromosome *mec* (SCC*mec*), carrying the methicillin-resistance gene *mecA*. SCC*mec* is currently classified into 13 types, with types I-V (20-60 kb) being the most prevalent and widely distributed (6). The cassette chromosome recombinases (Ccr) encoded by SCC*mec* (*ccrAB* for types I-IV, *ccrC* for type V) mediate its excision and integration at a specific attachment site (attB) located at the 3’end of the *orfX* gene (*rmlH*) in *S. aureus* chromosome (7, 8). The origins of SCC*mec* are not fully clear, but ancestral fragments have been identified in coagulase-negative staphylococci (CoNS) species and *Macrococcus caseolyticus* (6), and at least 20 independent acquisitions of SCC*mec* have been predicted to occur in *S. aureus* (9). Despite evidence of dissemination by horizontal gene transfer (HGT), the major mechanism of SCC*mec* transmission has remained a mystery for several decades (10).

Bacteriophage-mediated transduction and conjugation have been suggested as possible mechanisms of SCC*mec* transfer, but only short or fragmented SCC*mec* could be transmitted by these mechanisms (11, 12). Natural competence for DNA transformation is another HGT mechanism enabling bacteria to incorporate extracellular DNA through the DNA-uptake machinery encoded by a series of competence genes (13). This process in Gram-positive bacteria has been extensively studied in *Bacillus subtilis* and *Streptococcus pneumonia*. While the DNA-uptake machinery is highly conserved, the signalling and conditions regulating competence development are diverse among species, and finding the *bona fide* transformation conditions for a given specific species or strain is challenging (14). In *B. subtilis*, the transcription factor ComK controls the expression of competence genes in a subpopulation. ComK is tightly regulated by environmental signals such as quorum sensing and nutrient limitation. In *S. pneumonia*, competence is activated by the SigX sigma factor (a.k.a. ComX) in the whole population upon induction by quorum sensing and stressors. In *S. aureus*, a subpopulation expresses competence genes when grown in a complete synthetic medium (CS2). The competence operons, *comG* and *comE* operons, are under the transcriptional control of SigH and ComK (15–17). Recently, we showed that growth in biofilm conditions promotes natural transformation in *S. aureus* cells (18). In these biofilm conditions, we observed cell-to-cell inter- and intraspecies transfer of SCC*mec* by natural transformation. However, the transformation frequencies of SCC*mec* were low (∼10^-8^-10^-7^).

In this study, we aimed to seek the better biofilm conditions required for the SCC*mec* transformation. Our results show that mixed biofilms with living donor cells can be the appropriate place for SCC*mec* as well as other genetic elements to be transferred efficiently.

## Results

### Living donor cells enhance transformation efficiency of SCC*mec* in biofilms

To explore better biofilm conditions for SCC*mec* transfer, we tested different culture media, temperatures, donor cell status (heat-killed or alive), and the ratio of the donor to recipient (Fig. S1, Fig. 1). The laboratory strain Nef (N315 derivative that lacks conjugative genes, a lysogenic phage, and SCC*mec*) was grown statically (biofilm growth conditions) with donor cells for 3 days in one medium: CS2 medium (15, 18), diluted TSB (50% TSB), diluted RPMI (50% RPMI), or SMM (19, 20) at different temperatures: 25°C, 37°C, and 42°C. COLw/oφ (COL derivative that lacks conjugative genes and the lysogenized phage but carries SCC*mec*) was used as the donor. In Fig. S1, we used heat-killed donor cells as in previous reports (15, 18), while in Fig. 1, living donor cells were tested. In all experiments, a fixed amount of donor cells (∼5x10^8^ cfu per well equivalent) was used, while the recipient cell number was changed, generating the series of donor-to-recipient ratios shown in each figure. SCC*mec*-transformants were selected using the β-lactam antibiotic cefmetazole. Erythromycin was added to eliminate the living donor (Fig. 1). The ability of emerged colonies to grow in the presence of cefmetazole was confirmed by replica (18). The transformation frequencies were calculated as the ratio of the number of emerged transformants to the total cfu of the recipient at the end of the transformation assay and are shown in Fig. 1.

**Figure 1.**
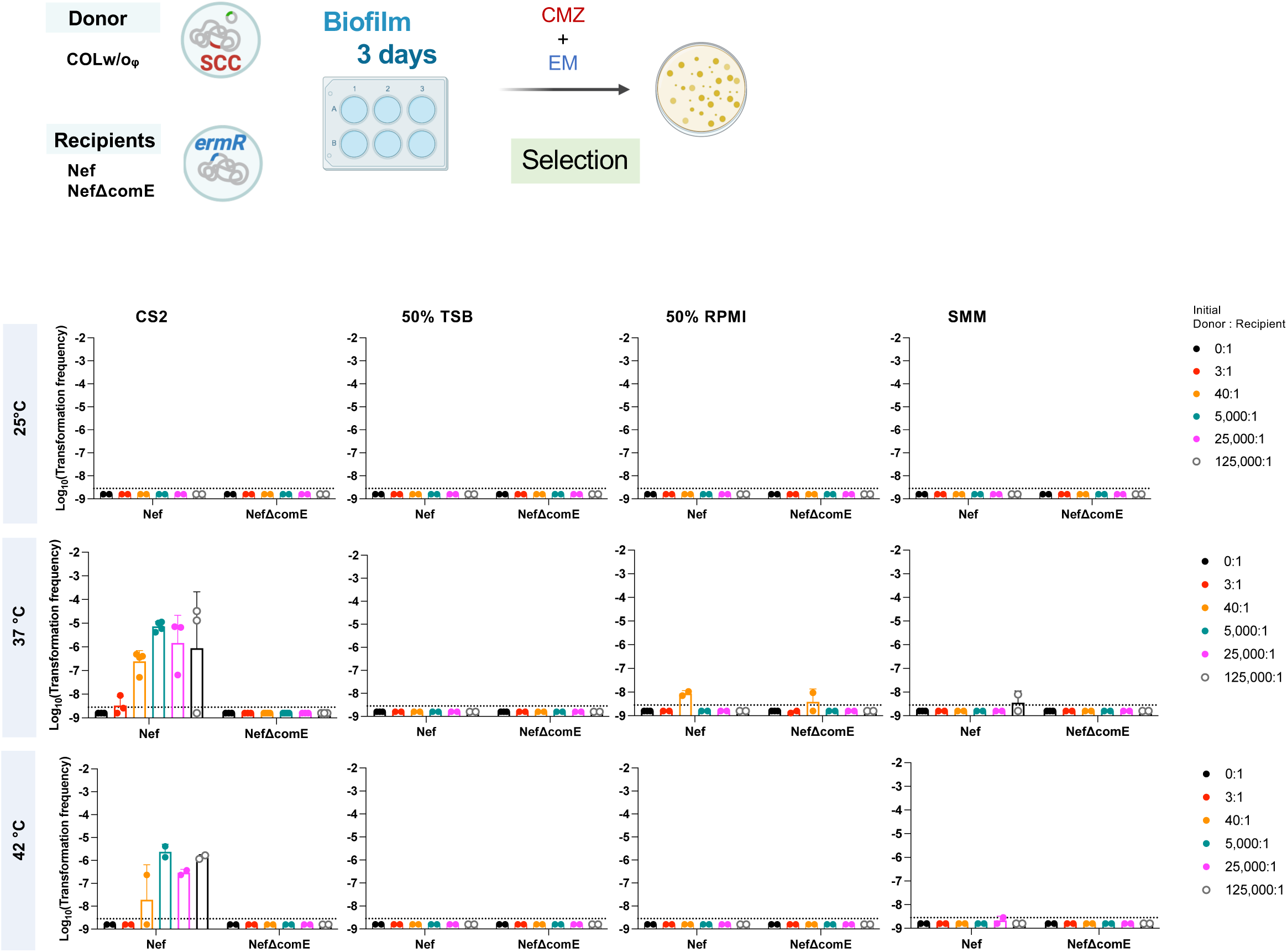
Transformation efficiency of SCC*mec* in the biofilms in different conditions using living donor. The recipient cells (Nef or NefΔcomE) were statically grown in different media and temperatures for 3 days. An initial fixed amount of living COLw/oφ donor cells was used, generating the following donor-to-recipient ratios 0:1, 3:1, 40:1, 5000:1, 25000:1, and 125000:1. Transformation efficiencies were determined after 3 days by selection with cefmetazole and erythromycin (EM). The mean of at least n = 2 independent experiments is shown with SD. Data points represent independent experiments. The dotted lines represent the detection limit.

The condition of our previously reported protocol using heat-killed donor is marked by the red arrow (Fig. S1: CS2, 37°C, 4:1), and the frequency was about 10^-8^. This culture condition (CS2 at 37°C) was one of the best experimental settings for transformation that we have found in previous studies, and this study again indicates that no other combination of the tested culture medium and temperature is better for transformation (Fig. S1). Increasing the ratio of the heat-killed donor to the recipient (reducing the number of recipient cells) did not improve the transformation frequency. Strikingly, on the other hand, when we used living donor cells instead of heat-killed cells, increasing the donor-to-recipient ratio resulted in higher transformation frequencies of SCC*mec* (up to 10^-5^∼10^-4^) (Fig. 1: CS2, 37°C). This efficient transfer of SCC*mec* from living cells is via natural transformation because no transformants were generated when NefΔcomE was used as the recipient. In a 50% RPMI medium, three cefmetazole-resistant colonies were detected from NefΔcomE recipient in one experiment, but it was not reproducible, and the reason is not known.

Time course of transformation using the living donor cells showed the transformation frequencies of SCC*mec* increasing towards day 2, with a peak around 10^-4^ on average and the maximum being 10^-3^-10^-2^, while the CFU values of recipient cells were sustained (Fig. 2A, see 5,000-125,000:1). This generated up to over 10^4^ transformants (5,234 on average) per one-well, including 10^7^ CFU recipient (Table S1, Fig. S3A). The generated transformants had the same genetic backbone as the recipient (Fig. S3) and full-length SCC*mec* (Fig. 1), as confirmed by PCR. These frequencies are drastically (10^4^-fold) higher than the frequencies reported in our previous study (18).

**Figure 2.**
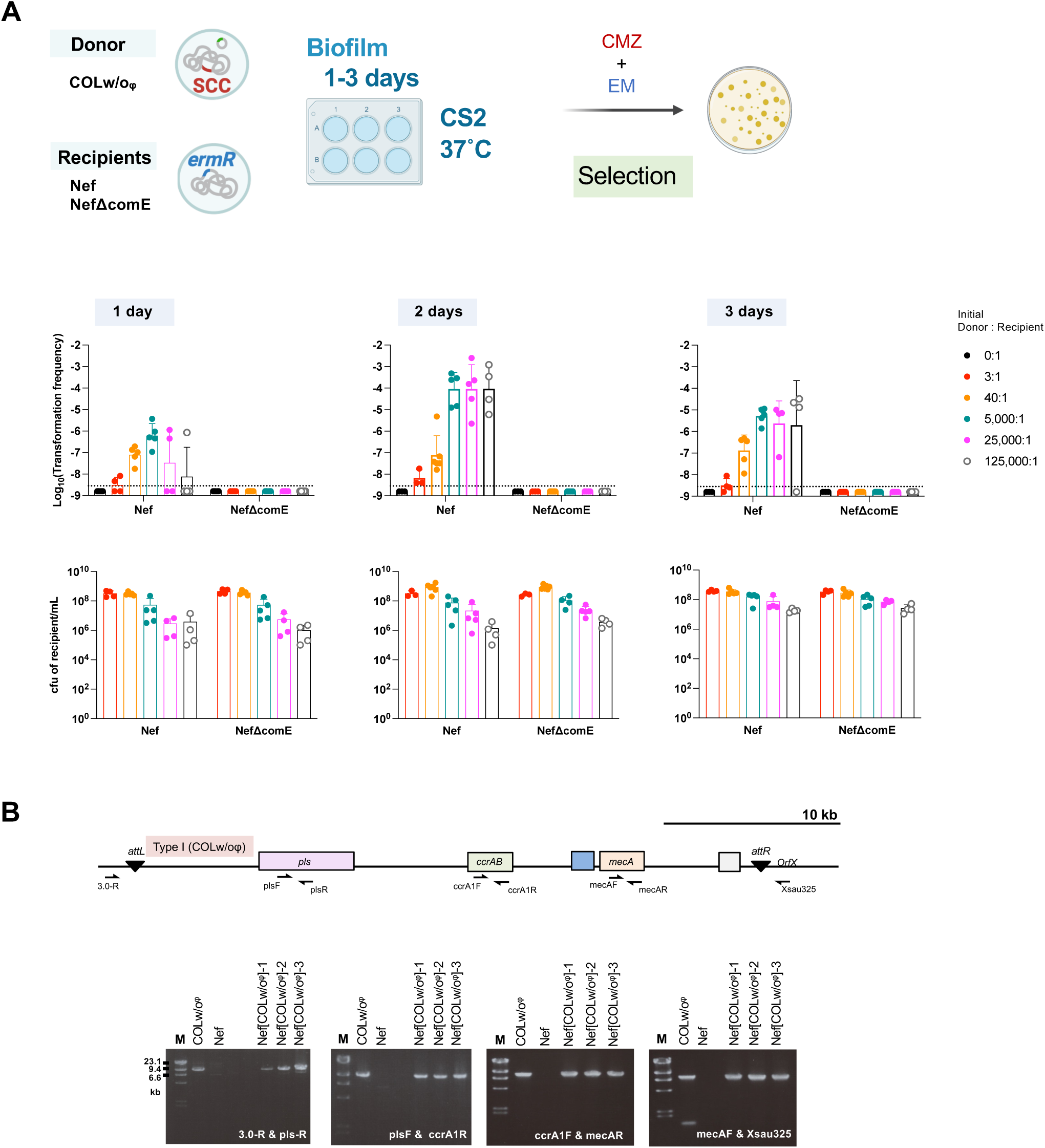
High donor-to-recipient ratio in 2-day biofilms enhances transformation efficiency of SCC*mec*. (A) The recipient cells (Nef or NefΔcomE) were statically grown in CS2 at 37°C for up to 3 days. An initial fixed amount of the living COLw/oφ donor cells was used, generating the following donor to recipient ratios 0:1, 3:1, 40:1, 5000:1, 25000:1, and 125000:1. Transformation efficiencies (top) or cfu/mL (bottom) of recipient cells were determined after every 24 h. Transformants were selected by cefmetazole (CMZ) and erythromycin (EM). The mean of at least n = 3 independent experiments is shown with SD. Data points represent independent experiments. The dotted lines represent the detection limit. (B) SCC*mec* I amplification in the Nef transformants obtained from 2-day biofilm using 5,000:1 donor-to-recipient ratio. The schematic structure of the SCC and primer locations indicated by arrows are shown. The DNA of COLw/oφ donor and Nef recipient was used for positive and negative controls. Suffixes (1), (2), (3) represent transformants obtained from three independent experiments. M: DNA marker, λ-HindIII.

### The Ccr-attB system is important for SCC*mec* transfer

SCC*mec* carries the *ccr* genes which encode recombinases that allow its excision and integration at the attB attachment site in the recipient’s chromosome. We tested whether SCC*mec* transfer is mediated by the Ccr-attB excision/integration system in the optimized transformation conditions using the living donor. Nef derivative strain that has a mutated attB site (attB*) showed a significant reduction (∼22-fold) in SCC*mec* transformation efficiency compared with Nef (Figs. S2, 3A). Moreover, when the living donor lacking the *ccr* genes (COLw/oφ-Δccr) was used, the transformation frequency was reduced ∼29-fold compared with the parental donor strain COLw/oφ (P=0.05) (Fig. 3A). Transformation frequencies of pT181 plasmid did not differ (Fig. 3B), suggesting that mutations in *ccr* and *attB* did not affect the efficiency of natural transformation itself.

**Figure 3.**
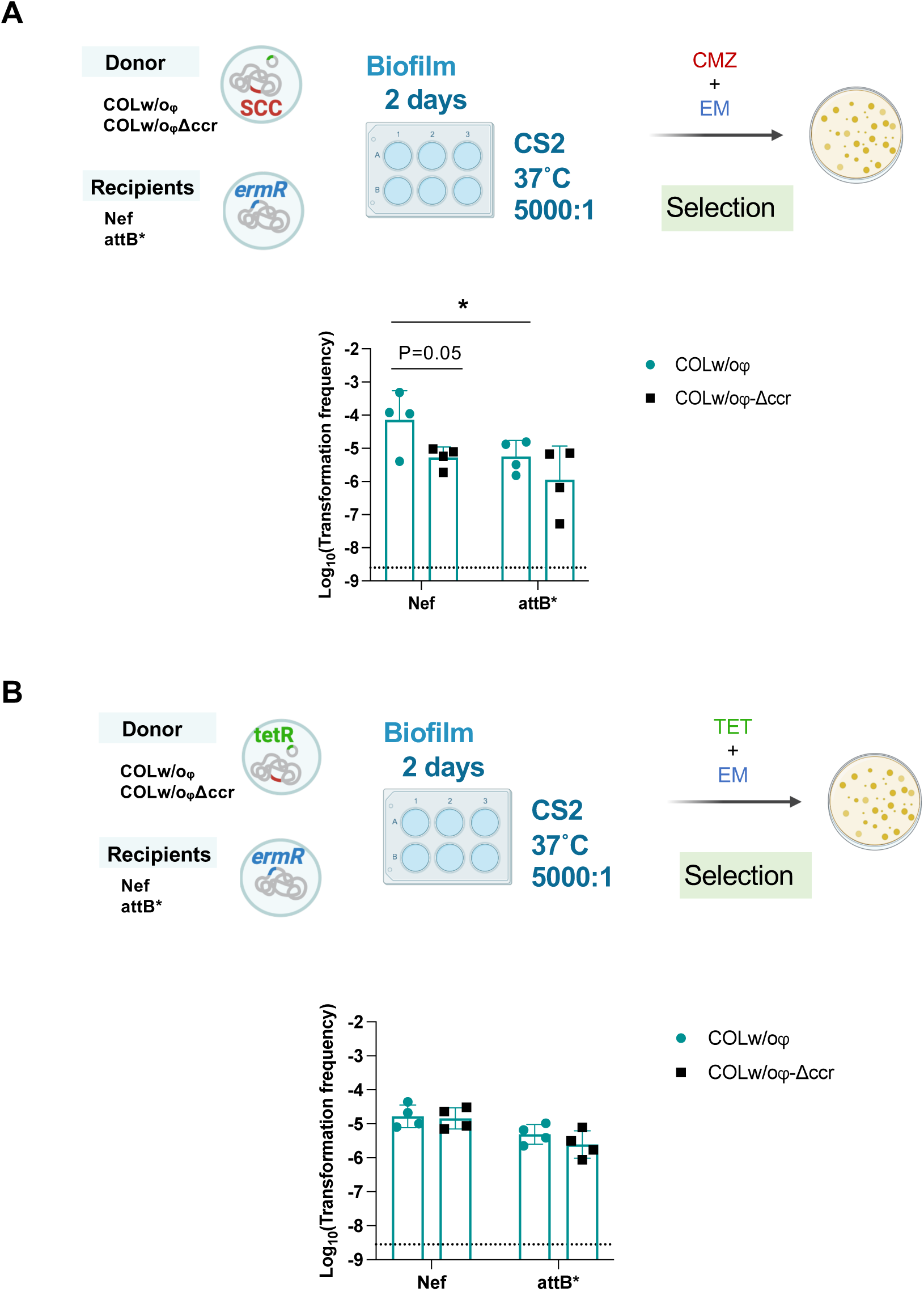
The Ccr-*attB* system is important for SCC*mec* transformation. (**A-B**) Living donor (COLw/oφ or COLw/oφ-Δccr) cells were added to recipient cells (Nef or attB*) in a 5000:1 ratio. The cells were statically grown in CS2 for 2 days at 37°C. Transformants were selected by erythromycin (EM) and cefmetazole (CMZ) (**A**), or by erythromycin and tetracycline (TET) (**B**). The mean of at least n = 3 independent experiments is shown with SD. Data points represent independent experiments. The dotted lines represent the detection limit. Statistical significance was determined by 2-way ANOVA followed by Tuckey’s multiple comparison tests as in Fig S2. *P<0.05.

### SCC*mec* can be transferred to clinical MSSA isolates in mixed biofilms

Previously, we showed that SCC*mec* can be transferred by natural transformation to clinical MSSA isolates in the biofilm using heat-killed donors (18). We aimed to test whether the transformation using living donors can more efficiently transfer SCC*mec* to clinical MSSA isolates. We used 10 erythromycin-resistant MSSA (98s, 9s-ermR, E1-E8) and erythromycin-sensitive MRSA/MRCoNS donors (COLw/oφ, MW2, MR-CoNS4, 11, 18), to use erythromycin to eliminate donors after transformation (Fig. 4). Nef, NefΔcomE, and 9sΔcomE-ermR strains were used as positive or negative controls. Five of the 10 tested clinical MSSA strains could generate SCC*mec*-transformants, and some of them showed high frequencies up to 10^-5^ (Fig. 4), which is much higher than in our previous report (∼10^-8^) (18).

**Figure 4.**
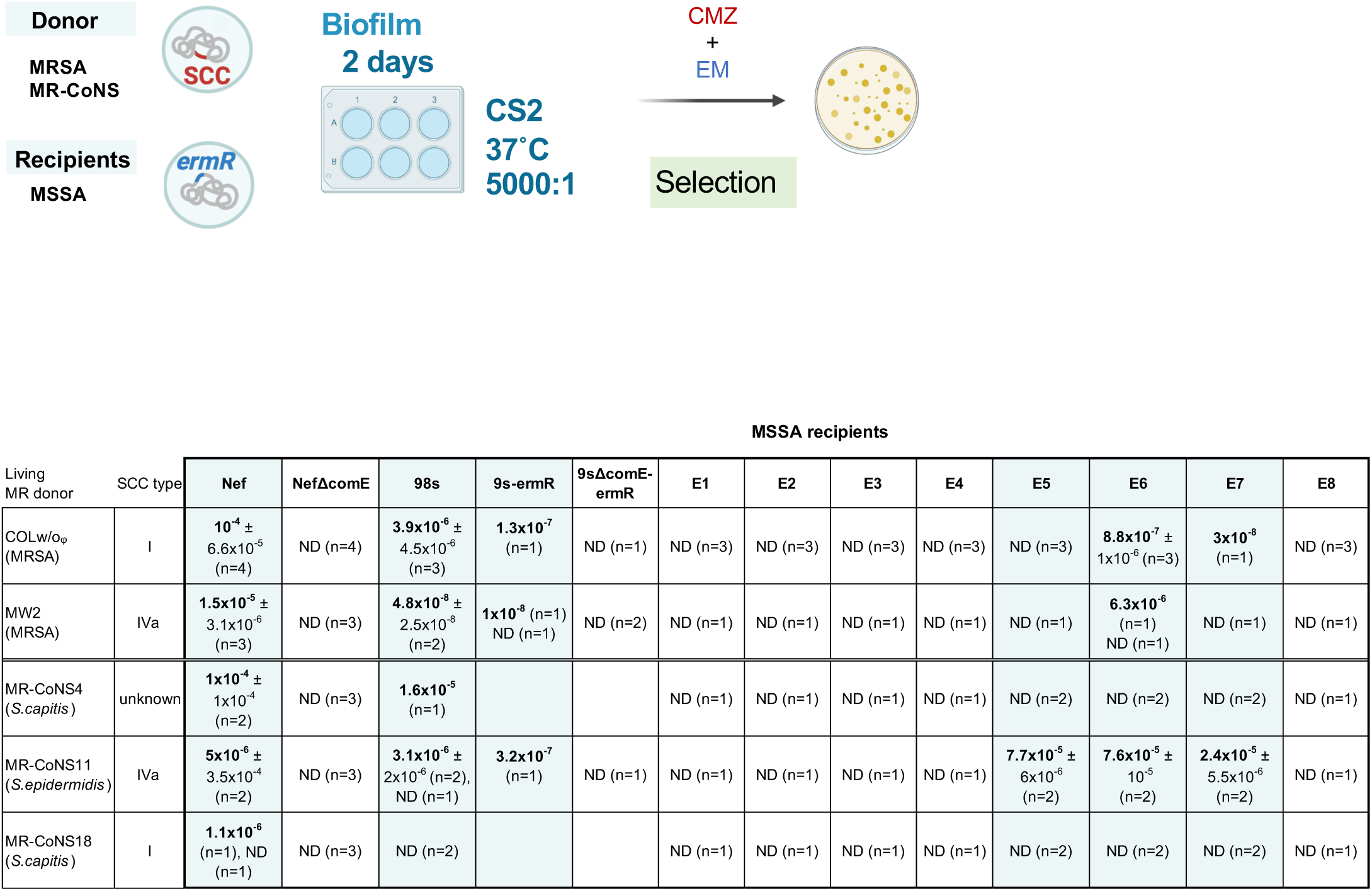
SCC*mec* can be transferred from living MR-CoNS donors in the biofilm. A fixed amount of living donor (MRSA or MR-CoNS) cells was added to different amounts of MSSA recipient cells in a 5000:1 ratio. The cells were statically grown in CS2 for 2 days at 37°C. Transformants were selected by erythromycin (EM) and cefmetazole (CMZ). The mean of n independent experiments ± SD are shown. ND, none-detected.

### Transformation of plasmid and chromosomal genes are enhanced by using living donor in the biofilm

In the optimized mixed biofilm conditions, we observed the transformation of *mecA* in the absence of the Ccr system (Fig. 3A). This implicates that the efficiency of the natural transformation of chromosomal genes would also be increased. Indeed, when we looked at the transformation frequencies of another chromosomal marker (the chloramphenicol resistance gene *cmR* in the donor COLw/oφΔcls1-cm^R^ strain), it was about 10^-4^-10^-3^ in the mixed biofilm conditions, while its transfer was undetectable in the previous protocol using heat-killed donors. Similarly, the transformation of plasmid pT181 (that has a tetracycline-resistance gene) was only detectable in the living donor system and about 10^-6^-10^-5^.

## Discussion

SCC*mec* is a large mobile genetic element conveying methicillin resistance in *Staphylococcus* and *Macrococcus* species (6, 21). The mechanism by which SCC*mec* transfers horizontally among staphylococci has been extensively researched in the past few decades in efforts to clarify how MRSA strains emerge and disseminate globally. Previously, we showed that natural transformation can mediate inter- and intraspecies transfer of SCC*mec* at low frequencies when the cells are grown in biofilm conditions (18). In this study, we present an enhanced transformation protocol that drastically improves the inter- and intraspecies transformation efficiency of SCC*mec* and other genetic elements in *S. aureus* biofilms. The high transformation frequencies detected in multiple strains and the dependency on the Ccr-attB system in SCC*mec* transfer support the importance of natural transformation in antibiotic resistance acquisition and MRSA emergence.

Previously, we reported that the heat-killed cells can serve as a donor in the biofilm transformation assays, while purified genomic DNA could not (18). In heat-killed cells, DNA resides within a peptidoglycan cell wall that is not destroyed by heat. It is conceivable that the cell wall structure reduces the DNA degradation by extracellular DNase (MNase or thermonuclease) that is known to be expressed in *S. aureus* biofilms (22). On the other hand, we confirmed that the heat-killed cells cannot completely block the DNase I access to the inside DNA, probably due to the destroyed membrane, while living cells can (18) (Fig. S15). The protection of donor DNA until its use by the competent cells would be one of the reasons for the present finding that living donor cells are better than heat-killed cells (Figs. 1, 5). In *B. subtilis*, the living donor cells have also been shown to enhance transformation efficiency and provide protection from DNase degradation (23). Another point to consider is that extracellular DNA sequestered in biofilm (24, 25) may not be able to serve as a donor for transformation, but it remains untested whether the DNA released from living donor cells with intact nucleoid proteins can serve as a good donor for transformation in biofilm.

This study did not address how the DNA is released from living donor cells and gets access to the recipient cells, but several mechanisms of DNA release have been suggested to occur in *S. aureus* biofilms, including autolysis, prophage-mediated lysis, vesicles, and active secretion systems (26–29). In our transformation assay, the phage-mediated lysis is dispensable, as the combination of COLw/oφ donor and Nef recipient does not include any phage components, but it remains a strong candidate in nature as most *S. aureus* isolates have at least one prophage. It might also be valuable to mention that growth in the CS2 medium causes unusual cell morphology (30), which may contribute to the DNA supply.

The enhanced transformation in this study requires a very high initial donor-to-recipient ratio (5,000:1 or more). This was not observed in *B. subtilis* transformation (23, 31), and cannot be explained simply by the importance of cell-to-cell contact or proximity between donor and recipient cells. The expression of *ccr* genes is limited to a subpopulation (32), and this might explain the necessity of a high donor amount for SCC*mec* transformation. However, the transfer of other genetic elements (plasmids and chromosomal genes) that are not dependent on *ccr* was efficiently enhanced as well (Fig. 5). One speculation is that a part of the donor cells continuously undergoes lysis, releasing the transforming DNA. Donor cells may also serve as a nutrient source for the growing recipient in the biofilm.

**Figure 5.**
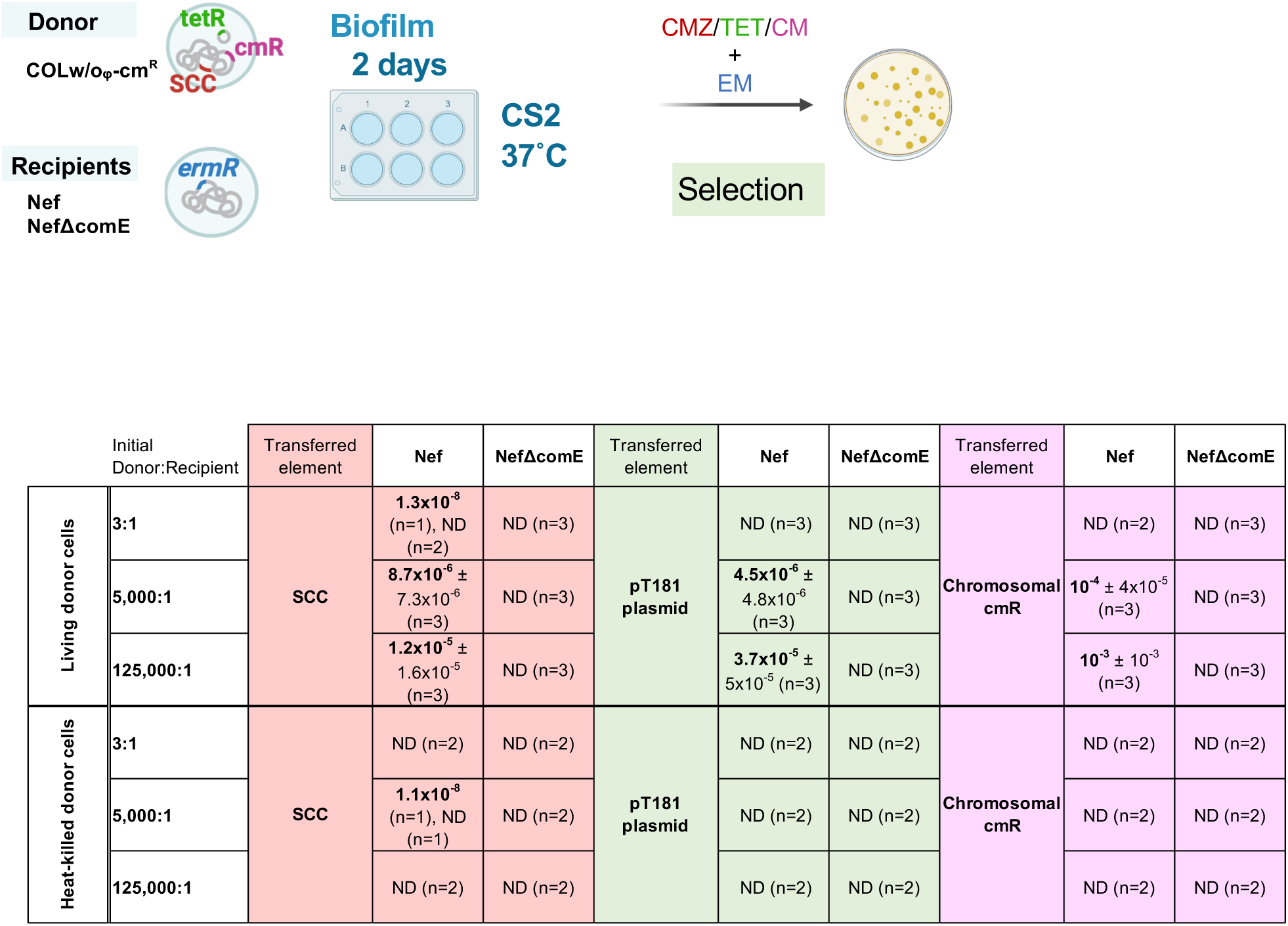
Living donor enhances transformation frequencies of genetic elements. A fixed amount of either living or heat-killed donor (COLw/oφΔcls1-cmR) cells was added to different amounts of recipient cells (Nef or NefΔcomE), generating the 3:1, 5000:1, or 125000:1 ratios. The cells were statically grown in CS2 for 2 days at 37°C. Transformants were selected by erythromycin (EM) with cefmetazole (CMZ) for SCC, or with tetracycline (TET) for the pT181 plasmid, or chloramphenicol (CM) for the chromosomal *cmR* gene. The mean of n independent experiments ± SD are shown. ND, none-detected.

The Ccr recombinases enable the excision and integration of SCC*mec* at the attB site in the recipient’s chromosome (7). The excision of SCC*mec* happens in a subpopulation that may later serve as a donor for horizontal transmission (32). SCC*mec* transfer was reduced but not abolished in the absence of the *ccrAB* genes on the donor side or in the absence of attB on the recipient side (Fig. 3). This minor Ccr-independent transfer would be through homologous recombination, as is observed for other chromosomal genes (Fig. 5). The importance of Ccr in the transformation observed in this study emphasizes the relevance of the natural transformation in SCC transmission among the HGT mechanisms. Whether Ccr is involved in other HGT mechanisms, such as phage transduction, is still elusive.

Polymicrobial biofilms are thought to be a hotspot for exchanging genetic material through HGT (33), and it is conceivable that staphylococcal biofilms also enhance HGT (10, 18, 34). In the present study, we established that a mixed biofilm enables the efficient transfer of SCC*mec*, providing insights into the environment in which MRSA is born. Importantly, many of the antibiotic-resistant genes, including *mec* genes, are shared between animal and human *Staphylococcus* species (35). The transformation method using mixed biofilm established in this study would be a valuable tool to study drug resistance and virulence dissemination among *Staphylococcus* species, as well as the factors affecting the SCC*mec* stability in *S. aureus* (18). In addition, it would also help provide a better understanding of the interplay between natural transformation and the different HGT mechanisms that might have co-evolved in biofilms.

## Materials and Methods

### Bacterial strains, plasmids, primers, and media

The bacterial strains and plasmids used in this study are shown in Table S3. The primers used in this study are shown in Table S4. Clinical staphylococcal samples (9 MSSA isolates and 4 MR-CoNS isolates) were collected from the Kanto area of Japan.

Staphylococci were grown in TSB for routine cultures. For biofilm formation and transformation assays, *S. aureus* was grown in CS2 (complete synthetic medium), diluted TSB, diluted BHI, diluted RPMI 1640, or SMM. *E. coli* strains were grown in LB. Where required for selection, the medium was supplemented with chloramphenicol (12.5 μg/mL), tetracycline (5 μg/mL), cefmetazole (4 μg/mL), erythromycin (16 μg/mL), or ampicillin (for *E. coli*, 100 μg/mL).

### Genetic characterization of staphylococcal strains

Multiplex PCR was performed using the QIAGEN multiplex PCR kit. The primer set is based on the previously reported methods to determine the CC types (36), with the addition of primers for *gyrA* (18). The clonal complex information shown in Table S3 is based on the amplification pattern of the target genes (36). Long amplifications of SCC*mec* were performed using KOD One PCR master mix (TOYOBO).

### Construction of mutants

Deletion mutants were constructed by double-crossover homologous recombination using the pMADcat vector (37). Fragments flanking the upstream (primers A and B, Table S4) and downstream (primers C and D, Table S4) regions of the locus targeted for deletion were amplified by PCR using chromosomal DNA from COLw/oφ as a template. The PCR products (AB and CD fragments) were used as templates for the overlap extension PCR using primers A and D (Table S4). The product was cloned into the *Bam*H I–*Sal* I site of pMADcat to generate the vector for *ccrAB* deletion (pMADcat-ΔccrAB, Table S3). The plasmids were purified from *E. coli* and were introduced into COLw/oφ after passaging through RN4220. Mutants (chloramphenicol sensitive, β-galactosidase negative) were selected (37) and the absence of the target gene was confirmed by PCR using the primers E and F (Table S4). The vector for *cls1* deletion (pMADcat1155, Table S3) was used as previously described (37) to generate the COLw/oφΔcls1-cm^R^ strain.

9s-ermR and 9sΔcomE-ermR strains were generated by transduction of the erythromycin-resistance gene from the Nef strain.

### Heat-killed donor preparation for natural transformation assays

Log-phase heat-killed donor cells were prepared as previously described (18). Overnight cultures were diluted 20-folds in TSB and were grown for 3 h at 37°C with shaking. Cells were then harvested in PBS (∼10^9^ CFU/mL) and boiled for 10 min. The absence of viable cells was confirmed by plating on TSB plates. For the SCC*mec* donor, COLw/oφ was used.

### Natural transformation assays using heat-killed donor

Natural transformation assays were conducted in a polystyrene 6-well plate under biofilm growth conditions (18). Briefly, 250 μL of log-phase heat-killed donor cells were added to 750 μL of log-phase recipient cells (∼10^8^ CFU/mL), and the total volume of growth medium was adjusted to 1.5 mL per well. To generate increasing donor-to-recipient ratios as indicated in Fig. S1, a fixed amount of donor cells (∼2.5 × 10^8^ CFU) was added to different amounts of recipient cells in a total volume of 1.5 mL pre well. The 6-well plates were incubated statically for 3 days at 37 °C and the medium was refreshed every 24 h. The biofilms were collected by pipetting and poured into BHI agar supplemented with cefmetazole (for SCC*mec* donors) for the selection of transformants. NefΔcomE strain was used as the negative control.

### Natural transformation assays using living donor

To detect natural transformation using living donors in the biofilm, 75 µL of donor overnight culture (∼10^9^ CFU/mL) were washed in the appropriate medium and added to different amounts of recipient cells from overnight cultures, that were washed in the appropriate medium, generating the ratios indicated in Figures 1-5. The cells were added to the polystyrene 6-well plate and the total volume of the growth medium was adjusted to 1.5 mL per well. The plate was incubated for the appropriate time and the medium was refreshed every 24 h. The collected biofilms were poured into melted BHI agar (precooled to 50°C) supplemented with appropriate antibiotics and solidified at room temperature. High concentration of erythromycin (16 μg/mL) was used to select cells derived from the erythromycin-resistant recipient, and this concentration can prohibit the growth of spontaneous mutants. The *mecA-*, *tetR-*, and *cmR*-transformants were selected by cefmetazole (4 μg/mL), tetracycline (5 μg/mL), and chloramphenicol (12.5 μg/mL). The transformants were confirmed by their ability to grow on replica and were verified by PCR (some examples are shown in Fig. S3). NefΔcomE strain was used as the negative control. In our experience, PCR test is essential for the chloramphenicol selection to eliminate spontaneous mutants, while we have not experienced spontaneous mutants for cefmetazole and tetracycline. In the case of *mecA* transformants, it is known that some are unstable depending on the combination of donor and recipient (18), but this study did not distinguish them to calculate the transformation frequency.

### Calculation of transformation frequency

Transformation frequency was calculated as the ratio of the number of transformants to the total CFU of the recipient after transformation. None detected values were assigned half the value of the detection limit for the calculation of mean values and statistical analyses.

### Statistics

Statistical analyses were performed by GraphPad Prism (GraphPad Software, version 8.4.3) on data from three or more independent experiments. Error bars indicate SD. The difference among groups was analyzed by two-way ANOVA followed by Tukey’s multiple-comparison test, as indicated in figure legends. The log values of natural transformation frequencies were analyzed statistically. \**P* < 0.05 was considered statistically significant.

## Acknowledgements

We thank our laboratory members for the discussions. This research was supported by AMED Grant Number JP23fk0108630, JSPS Bilateral Program Grant Number JPJSBP120229908, JSPS KAKENHI Grant Number 22H02863 (to KM), JSPS KAKENHI Grant Number 23K14515 (to MM).

## Author contributions

K.M. conceived the study and supervised the research. M.M. performed experiments and interpreted data with K.M. and Y.U. M.H. contributed to the study design. M.M. and K.M. wrote the paper. All authors checked and approved the final manuscript.

## Supplementary information captions

**Figure S1. Transformation efficiency of SCC*mec* in the biofilms in different conditions using heat-killed donor.**

The recipient cells (Nef or NefΔcomE) were statically grown in different media and temperatures for 3 days. An initial fixed amount of the heat-killed COLw/oφ donor cells was used, generating the following donor to recipient ratios 0:1, 3:1, 40:1, 5000:1, 25000:1, and 125000:1. Transformation efficiencies were determined after 3 days by cefmetazole (CMZ) selection.

The mean of at least n = 2 independent experiments is shown with SD. Data points represent independent experiments. The dotted lines represent the detection limit.

**Figure S2. The *attB site* is important for SCC*mec* transformation.**

Living donor COLw/oφ cells were added to recipient cells (Nef or attB*) in a 3:1, 5000:1, or 25000:1 ratio. The cells were statically grown in CS2 for 2 days at 37°C. Transformants were selected by erythromycin (EM) and cefmetazole (CMZ) The mean of at least n = 3 independent experiments is shown with SD. Data points represent independent experiments. The dotted lines represent the detection limit. Statistical significance was determined by 2-way ANOVA followed by Tuckey’s multiple comparison tests. *P<0.05.

**Figure S3. Confirmation of transformants by PCR.**

(A) Representative photo of SCC*mec*-transformants using COLw/oφ as donor and Nef as the recipient in a 5000:1 ratio, from 2-day biofilm.

(B) Transformants share the same genetic background as the recipient, as validated by multiplex PCR: The amplification patterns are the same between transformants and each original recipient. M: 100 bp DNA ladder (Takara).

(C) The SCC*mec*-transformants have the *mecA* gene. Genomic DNA of the donor and recipient was used as positive and negative controls respectively.

(D) Transformants carry the pT181 plasmid. Genomic DNA of the donor and recipient was used as positive and negative controls respectively. M: λ-HindIII DNA ladder (Takara).

(E) SCC*mec* IVa amplification in the Nef transformants obtained from 2-day biofilm using a 5,000:1 donor-to-recipient ratio. The schematic structure of the SCC and primer locations indicated by arrows are shown. DNA of COLw/oφ donor and Nef recipient was used for positive and negative controls. Suffixes (1), (2), and (3) represent transformants obtained from three independent experiments. M: DNA marker, λ-HindIII.

**Table S1. The number of transformants in biofilm obtained from living COLw/o**φ **donor cells (from** Fig. 2).

**Table S2. The number of transformants in 2-day biofilm obtained from living COLw/oφΔcls1-cm^R^ donor cells.**

**Table S3. Bacterial strains and plasmids used in this study. Table S4. List of primers used in this study.**

